# Ecological strategy and genetic load in the shepherd’s purse (*Capsella bursa-pastoris*) from the core and the limit of its natural range

**DOI:** 10.1101/563569

**Authors:** Marion Orsucci, Pascal Milesi, Johanna Hansen, Johanna Girodolle, Sylvain Glémin, Martin Lascoux

## Abstract

Species range expansion is a complex process whose outcome depends on the interplay of demographic, environmental and genetic factors. In plants, self-fertilizing species that do not require a mate to reproduce usually show higher invasive ability. However, this comes at a cost as both selfing and bottlenecks occurring during colonization lead to an increase in deleterious mutations accumulation (genetic load). Although they are theoretically clearly spelled out, the relationships between genomic and phenotypic characteristics of expanding populations have hitherto rarely been characterized.

In the present study we analyzed how different accessions of the shepherd’s purse, *C. bursa-pastoris*, coming from the front of colonization or from the core of the natural range performed under increasing density of competitors. We first showed that, as expected, accessions from the front of colonization performed the worst for most life history traits compared with accessions from core populations. Second, competitor density had a negative impact on both vegetative growth and reproductive output in term of fruits production for all accessions. However, somewhat unexpectedly given their higher genetic load and their lower absolute performance, accessions from the front of colonization were less affected by competition than accessions from the core of the species range. This could be due to phenotypic tradeoffs and a shift in phenology that allow the accessions from the front of colonization to avoid competition. These results are discussed in terms of ecological strategies of expanding populations.

## Introduction

Species range expansion is a complex process whose outcome depends on interplay between demographic, environmental and genetic factors. High dispersal ability is obviously key to reach new unoccupied habitats but colonization success also depends on other ecological processes. While initial establishment can be favored because within species competition is initially low, this low individual density also means low mate availability. By consequence, in plant species, the ability to self-fertilize is expected to increase colonization ability (Pannell 2015) and indeed self-compatibility and selfing are often associated to weedy habit (Clements et al. 2004), invasiveness (Van Kleunen & Johnson 2007; Van Kleunen et al. 2010) and ruderal strategies (Munoz et al. 2016). In pairwise comparisons, selfing species have also been shown to have larger species range than their outcrossing relatives (Grossenbacher et al. 2015). However, range expansion and colonization of new habitats can eventually be costly. First, a colonization/competition trade-off is expected such that a good colonizer can be a poor competitor (Burton et al. 2010), which can limit range colonization to sufficiently open and disturbed habitats. Second, the peculiar demographic dynamics associated with range expansion, namely, recurrent bottlenecks followed by rapid population growth, can lead to a genetic process known as “allele surfing” (Excoffier et al. 2009), which can contribute to the accumulation of deleterious mutations on the expansion front, creating a so-called “expansion load” (Peischl et al. 2013; Peischl & Excoffier 2015; Willi et al. 2018). Alternatively, ecological strategies to avoid competition, such as phenological changes can evolve within newly colonized populations. Plant phenology varies over space and time thereby allowing species coexistence by using alternative resources or using the resources at different moments (Chesson 2000), and early flowering can be favoured in newly colonized populations if early flowering plants face less competition.

In addition to selfing, allopolyploidization, which increases genome copy number through species hybridization, can also confer an immediate advantage when colonizing new habitats. Both mechanisms could explain the colonization and invasion successes of several plant species: self-fertilization provides reproductive insurance, a single individual being able to initiate an invasion (Barrett & Husband 1990; Hollingsworth & Bailey 2002; Pettengill et al. 2016; Munoz et al. 2016) while polyploidy allows a partial sheltering from the negative effect of selfing, especially by masking deleterious alleles (Hahn et al. 2012; Beest et al. 2012).

Although they are theoretically clearly spelled out, the relationships between genomic and phenotypic characteristics of expanding populations have hitherto rarely been characterized. Direct characterization of fitness and accumulation of mutations during range expansion have only been performed in experimentally evolving populations of bacteria (Bosshard et al. 2017) or in North American populations of the plant species *Arabidopsis lyrata* that recently went through a range expansion (Willi et al. 2018). In *A. lyrata*, only the global performance of individual plants was measured so we still do not know whether and how different parts of the life cycle and individual competitive ability were affected during range expansion.

The shepherd’s purse, *Capsella bursa-pastoris* (L.) Medik. (Brassicaceae), a species with a very large distribution due to a recent range expansion (Cornille et al. 2016)), is a good model species to characterize the evolution of the life cycle and the possible effect of deleterious mutations during range expansion. It is an annual, selfing and allotetraploid species originating from the hybridization, ca. 100 to 300 k years ago, of the ancestral lineages of two diploid species of the same genus, the obligate outcrosser, *C. grandiflora* (Klokov) and the self-fertilizing *C. orientalis* (Fauché & Chaub.) Boiss. (Douglas et al., 2015; Kryvokhyzha et al. 2019). *C. bursa-pastoris* probably first occurred in the Middle East (ME), then spread to Europe (EUR) before invading Eastern Asia (ASI) and more recently spread worldwide due to human migrations (Cornille et al., 2016). Due to limited gene flow between these geographic areas, three main genetic clusters can be delineated from both nuclear and expression data (Cornille et al. 2016; Kryvokhyzha et al. 2016; Kryvokhyzha et al. 2019). Furthermore, as expected given the species history the ASI populations (*i.e*. colonization front) presented a reduced genetic diversity and a higher amount of deleterious mutations than the EUR and ME populations (core populations; Kryvokhyzha et al. 2019).

Two previous studies have started to investigate the differences in competitive ability among *Capsella* species and showed that (i) selfing species of the *Capsella* genus were more sensitive to competition than outcrossing ones (Petrone Mendoza et al. 2018; Yang et al. 2018) and (ii) competitive ability decreased during range expansion in *C. bursa-pastoris* (Yang et al. 2018). However, these studies did not consider all stages and fitness components of the life cycle and did not directly test for a possible effect of the load accumulated during range expansion.

Here, we conducted an experiment under controlled environment following the whole life cycle of *C. bursa-pastoris* accessions whose number of potentially deleterious mutations had been estimated from whole genome sequence data (Kryvokhyzha et al. 2019). Life history traits, such as fitness components (fertility, viability of the progeny), phenological traits (flowering start, germination time) and vegetative traits (growth of the rosette) were assessed in *C. bursa-pastoris* accessions coming from the three main genetic clusters. In contrast with most studies based on one (or few) life-history traits, we recorded the impact of different density of competitors on traits associated with the complete life cycle, *i.e*. from the mother plant to the progeny. This allowed us measuring both fertility and viability, two major components of fitness, and to highlight the presence of trade-off between traits. We showed that competitor density impacted only the fertility of the mother plant but not its viability. More generally, we detected that populations from the colonization front (Asian accessions) had a lower fitness, i.e. less fruits and lower germination rate, than populations from the core of the distribution (European and Middle-Eastern accessions). However, contrary to our expectations, Asian accessions, which carry a larger number of deleterious mutations than European and Middle-Eastern ones, were the least affected by competition in the conditions of the experiment. We show that this could be due to their very early flowering start, which allows them to partially avoid the effect of competition. These results also highlight the need to take the whole life cycle into account to better infer fitness and understand trait evolution.

## Materials et methods

### Plants sampling and seeds preparation

#### Sampling

We assessed the competitive ability of accessions of *Capsella bursa-pastoris* (*Cbp*) from different parts of the natural range against different numbers of competitors. As competitor we chose an annual species growing in similar environments, *Matricaria chamomilla* (Asteraceae) (Fig. 1A). The two species have similar distribution areas (Europe and temperate Asia) and do co-occur, in particular in Northern Greece where *M. chamomilla* appears as one of the main competitors of *Capsella* species (S. Glémin and M. Lascoux, pers. observations). We used commercial seeds of *Matricaria chamomilla* in order to ensure good germination and homogeneity among plants. We used 24 accessions of *C. bursa-pastoris*, corresponding to seeds harvested on a single maternal plant originating from different sampling sites that belong to one of the three main genetic clusters, Europe, Middle East and Asia, as identified by Cornille et al. (2016) and Kryvokhyzha et al. (2016). More specifically, nine accessions came from sites distributed across Europe (EU), 11 from sites in Asia (AS), three from sites in the Middle East (ME) and one from the United-States (Fig. 1B; Table S1). The latter belongs to the ME genetic cluster and genomic and transcriptomic data (Kryvokhyzha et al. 2016) suggests a likely very recent introduction; it was thus added to the other ME accessions in following analyses.

**Figure 1.**
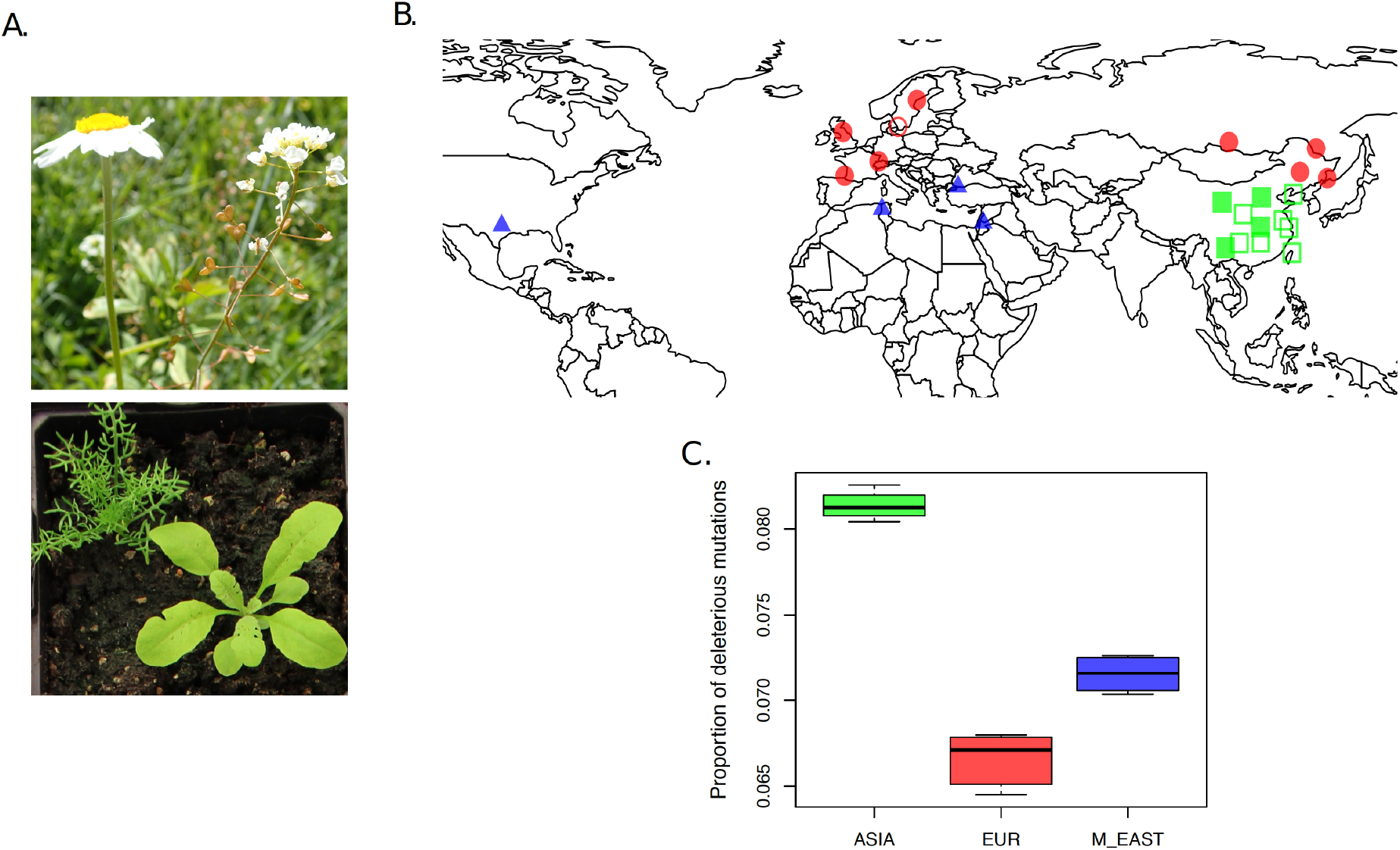
Capsella bursa-pastoris. **A.** *C. bursa-pastoris* and *Matricaria chamomilla* in natural conditions and in our experiment. **B.** Distribution of *C. bursa-pastoris* accessions used in the competition experiment (Europeans are red dots, Middle-Eastern are blue triangles and Asian accessions are green squares); empty symbol means that the germination rate was too low to include the accession in the experiment. **C.** Proportion of deleterious mutations carried by the different accession from Asia (AS, green), Middle-East (ME, blue) and Europe (EUR, red).

#### Seed preparation

For each accession, at least 30 seeds were surface-sterilized and sown into agar plates (Table 1). Following sterilization, seeds in agar plates were kept during 7 days at 4°C and in complete darkness for stratification. Agar plates were then transferred into a growth chamber (12:12h light:darkness cycles, 22°C) for germination (which starts approximately two days later). After one week, the seedlings that presented a radicle of at least 1 cm long and well-developed cotyledons were used for the competition experiment.

**Table 1.** *C. bursa-pastoris* germination rate.

| Origin area | Country | ID acc. | $GR_{mp}$ |
| --- | --- | --- | --- |
| Asia | China | DL174 | 0.05 |
|  | China | GY37 | 0 |
|  | China | HJC419 | 0 |
|  | China | HY85 | 0 |
|  | China | JZH152 | 0.64 |
|  | China | KMB206 | 0.98 |
|  | China | N444 | 0.91 |
|  | China | NJ219 | 0.08 |
|  | China | TBS195 | 0.07 |
|  | China | TY118 | 0.5 |
|  | Taiwan | PL | 0 |
| Europe | China* | HRB132 | 0.40 |
|  | France | FR50 | 0.93 |
|  | France | STJ2 | 0.24 |
|  | Russia | BEL5 | 1 |
|  | Russia | IRRU2 | 0.69 |
|  | Russia | VLA3 | 1 |
|  | Sweden | SE14 | 0.93 |
|  | Sweden | SE33 | 0 |
|  | UK | STA4 | 0.31 |
| Middle-East | Algeria | AL87 | 1 |
|  | Jordan | JO56 | 0.58 |
|  | Turkey | TR73 | 0.83 |
|  | USA | WAC5 | 0.68 |

For each geographic area, ID acc. is the accession identification and $GR_{mp}$ is the germination rate (minimum and average number of seeds sown per accession were respectively 33 and 47). Accessions that were not used in competition experiment because of a too low germination rate ( $GR_{mp} < 0.3$ ) are italicized. \*North China.

### Competition experiment

#### Experimental set-up

To assess the competitive abilities of the different *Cbp* accessions, we measured different life history traits in the absence (negative control) or in the presence of one, two, four or eight competitors (*Matricaria. chamomilla*). Each accession of *Cbp* was thus exposed to a total of five treatments (Fig. S1). The seedlings were transplanted in square pots of 11 x 11 cm and maintained in a growth chamber with 12:12h light:darkness cycles and a constant temperature of 22°C. This pot size allowed maximizing the competition while still permitting the focal plant development (Petrone Mendoza et al. 2018). The competitors (*M. chamomilla*) were transplanted three days before *Cpb* accessions because of their slightly slower development (data not shown). Then, for each *Cbp* accession, one individual was transplanted in the center of the pot surrounded by competitors (average distance between the focal plant and competitor(s) ~4cm, Fig. 1A and S1). All the treatments for each *Cbp* accession were replicated four times (*block*) and pots within each block were randomized. The complete design thus comprised 20 seedlings per accession (five competitive treatments x four blocks) and only accessions with a high enough germination rate (*GR_MP_* > 0.1) were used (Table 1).

#### Capsella bursa-pastoris accession performances

To assess viability and fertility of the accessions facing different competition intensities, several phenotypic traits were recorded. The traits were related to vegetative growth, phenology and fitness (Fig. S1). First, the diameter of the rosette of each focal plant (in cm) was measured 14 (Dt_1_) and 21 (Dt_2_) days after transplantation. The diameter of the rosette corresponds to the largest distance between opposite leaves (measured from pictures with the image analysis program, *Image J*, Schneider et al. 2012). The growth of rosette size (Δ_*growth*_) was then computed from Dt_2_ and Dt_1_ as Δ_*growth*_= Dt_2_ - Dt_1_. Second, we recorded three phenological traits corresponding to different life cycle stages: (*i*) the *flowering start* (*FS*) is the date of the first flower appearance (at least one white petal visible); (*ii*) the *flowering timespan* (*FT*) is the number of days between *FS* and plant death; (*iii*) the *lifetime* (*LT*) is the number of days from plant transplantation to death. Finally, two major components of fitness were assessed: (*i*) the *fertility, NF*, was estimated as the number of fruits produced (number of flowers and stems were also recorded, and were well correlated to *N_F_*), and (*ii*) the *germination rate of the progeny* either in soil (*GR_SOIL_*) or in agar (*GR_AGAR_*), see details below.

After the focal plant had dried out, up to 200 seeds were collected to investigate the fertility of the accession and the viability of its progeny. First, the *mean seed weight* (*W*), which is known to be a predictor of seed germination, was quantified as the weight of pooled seeds divided by the number of seeds in the pool (*ca*. 200 seeds in most case). Then, 25 randomly sampled seeds were sown on agar plate (*GR_AGAR_*) and 50 others directly in soil of square pots (7 x 7cm) to consider a more “natural” environment. After seven days of stratification (24h dark, 4°C), petri dishes and pots were moved into growth chambers (12:12h light:dark, 22°C). We studied the *germination dynamic* (*GD*) by recording the germination rate (number of germinated seeds over number of seeds sown) after one, two and seven days for the seeds in agar plates (*GD_AGAR_*) and after two, five, seven and fourteen days for seeds in pots (*GD_SOIL_*). The number of seed germinated over number of seeds sown at the last day of each treatment was considered as the progeny *germination rate* (*GR_AGAR_* and *GR_SOIL_*).

### Data analysis

#### Genetic load estimation

Kryvokhyzha et al. (2019) quantified the proportion of deleterious mutations that accumulated after the origin of *C. bursa-pastoris* carried by each subgenome in each accession. For each accession, we used the average proportion of deleterious mutations as an estimation of the genetic load (*del* = (*del_Co_* + *del_Cg_*) / 2).

#### Phenology

Cox’s hazard models (*coxph* function, *survival* package v.2.41-3, Therneau & Grambsh 2000; R software v.3.3.1, R core Team, 2016) were used to assess the effect of plant geographic origin (*G_OR_*, a three-level factor, AS, EU and ME) on flowering start (*FS* and *FS_prog._*) and progeny germination dynamics (*GD_AGAR_* and *GD_SOIL_*).

#### Phenotypic traits

For the phenotypic traits (vegetative and fitness-linked), generalized linear models (GLM, *glm* function, *stats* package v.3.6.0; R) were adjusted to the data to assess the effect of plant geographic origin, of the number of competitors and of their interaction, while controlling for block effect:

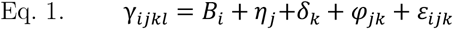

For a given trait, *γ* is the observation for a plant *l* from geographic origin *k*, in block *i*, and with *j* competitors. *B* is the block-specific fixed-effect (four-levels factor), *η* is the competition-specific fixed-effect, *δ* is the geographic origin fixed-effect (a three-level factor corresponding to the Asian, European and Middle-Eastern genetic clusters), *φ* is the interaction between factors *η* and *δ* and *ε* is the error term; the error term follows a binomial distribution for *GR_MP_, GR_AGAR_* and *GR_SOIL_*, a normal distribution for *W* and Δ_*growth*_ and a negative binomial distribution for *fertility* (*N_F_*). When the interaction term between the geographic origin and the number of competitors was significant, sub-models were used to estimate the effect of each factor independently.

When the geographic origin had a significant effect on a given trait, correlations between the trait and the average proportion of deleterious mutations between subgenomes (*μ_del_*) was estimated. As accessions from AS had larger values of *μ_del_* than the accessions from EU and ME, we reported Spearman’s rho, which is more conservative than Pearson’s product moment correlation if outliers are present. For comparison Spearman’s rho and Pearson’s product moment correlations are both reported in Table S2. Finally, to investigate phenotypic trade-offs, between-trait correlations were also estimated.

#### Competition index

To characterize the influence of competition intensity on the fitness of *Cbp* accessions, we defined a competition index (*I_C_*), as in Mendoza et al. (2018), based on a major component of fitness, the number of fruits produced (*N_F_*). For a given accession *k*, and *i* competitors (*i* = 1, 2, 4 or 8), *I_C_* was computed as:

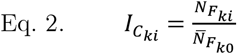

where 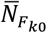 is the average number of fruits in absence of competition across the four replicates of accession *k*; the lower the *I_C_*, the higher the effect of competition. Significance of differences in response to competition for the various geographic origins was then assessed through GLM with a gamma error term, with block (*β*) and geographic origin (*δ*) as fixed-effects. Finally, for a given accession *k* the average competitive index was used as an indicator of the sensitivity to competition.

#### Model simplification

To assess the significance of the various fixed-effects, models were simplified by first removing higher order effects (interaction terms). The significance of difference in deviance explained between two models was assessed through a likelihood-ratio-test (Lrt). Significance of difference between factor-levels was tested through the same procedure and factor-levels that were not significantly different were merged (*p* > 0.05).

### Results

For each life history trait, a complete generalized linear model (GLM) that included the geographic origin (*G_OR_*), the number of competitors (*N_C_*), their interaction (*G_OR_* x *N_C_*) and a block effect (*B*) as explanatory variables was implemented. When the interaction, *G_OR_* x *N_C_*, explained a significant part of the total variance, sub-models were computed to estimate the effect of each factor independently. Globally, most of the life history traits differed among geographic areas (Fig. 2, Tab. S3 and Fig. S2). The first principal component explained 31% of the total variance and discriminated the three geographic areas, with the Asian cluster tending to be the more differentiated. The second principal component explained up to 21% of the total variance and mainly reflected the influence of competition intensity on life history traits. For clarity, we will first focus on the effect of geographic origin on each of the life history traits and then present the effects of the number of competitors.

**Figure 2.**
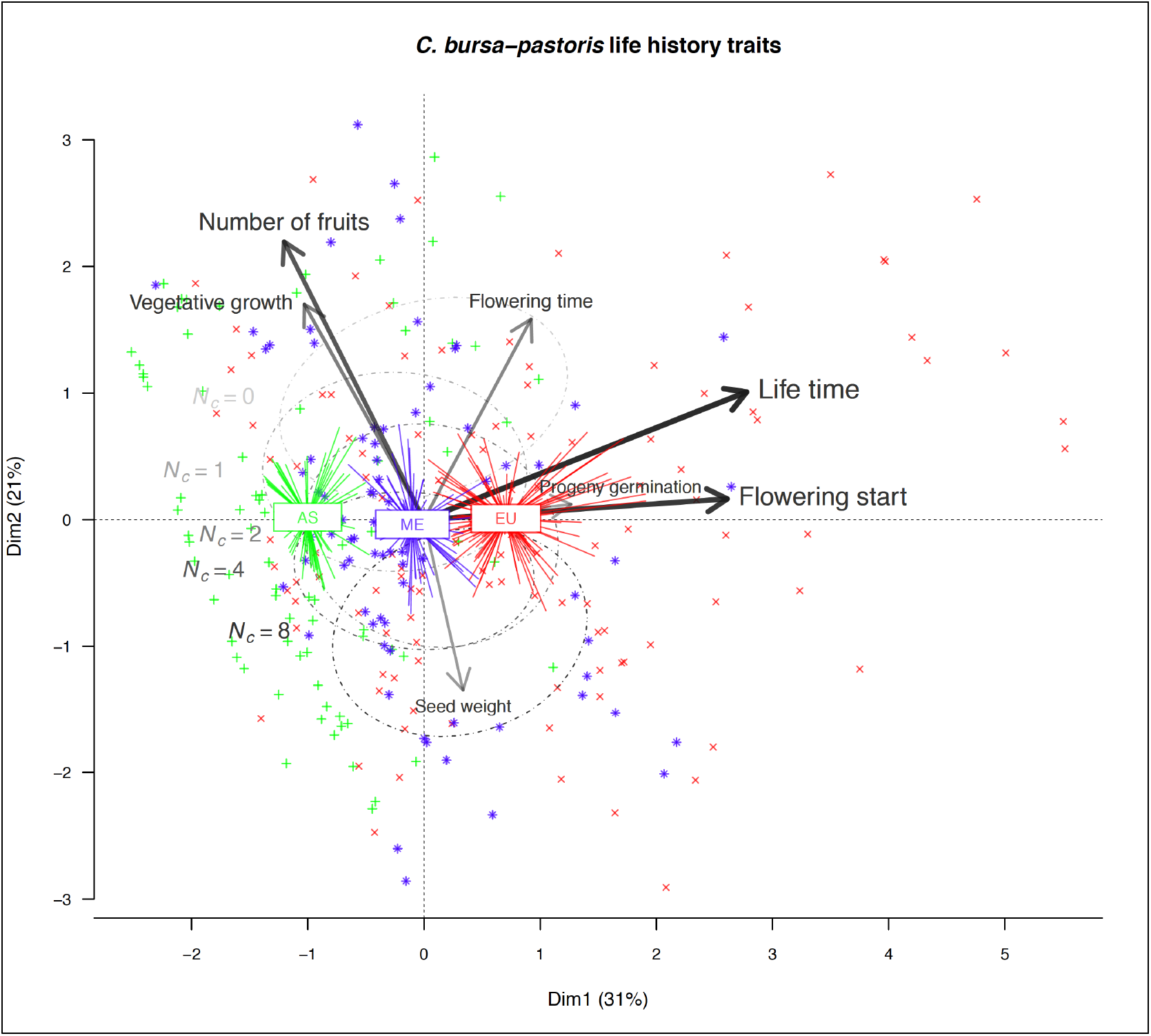
Principal components analysis based on *C. bursa-pastoris* life history traits. The first principal component (Dim1) mainly discriminates samples according to their geographic origin (Asia, AS green plus-signs; Middle-East, ME blue stars and Europe, EU red crosses) while Dim2 mainly reflects the intensity of competition (gray ellipses, *N_C_* = 0 to *N_C_* =8). Arrows represent phenotypic traits relative contribution, the thicker and darker the arrow, the higher the contribution.

### Accessions from the colonization front performed the worst for most life history traits but flowered early

#### Germination

The germination rate (*GR_MP_*) of Middle-Eastern accessions of *Capsella bursapastoris* (ME, 0.77 ± 0.1) was larger than that of European (EU, 0.61 ± 0.1) or Asian accessions (AS, 0.29 ± 0.1; Lrt, *Δdf* = 1, all *p* < 0.001; Fig. S2A). In addition, *GR_MP_* was negatively correlated to the amount of deleterious mutations of each subgenome, *i.e*. the higher the average number of deleterious mutations (μ_del_), the lower the germination rate (*ρ* = −0.49, S = 3418, *p* = 0.02). Finally, eight accessions (seven Asian and one European accessions) that had a germination rate < 0.3 were not included in the competition experiment (Table 1).

### Mother plants

#### Vegetative trait

Accessions from the Middle-East grew faster (Δ_*growth*_ = 1.2 ± 0.3 cm/days, Lrt, *χ*^2^ = 311,Δdf = 1,p < 0.001) than accessions from Europe (EU = 0.87 ± 0.06) or Asia (AS = 0.85 ± 0.12) that did not differ significantly (Lrt, *χ*^2^ = 0.5, Δ*df* = 1,p = 0.82, Table 2 and Fig. S2B1); however, this difference in growth rates was not correlated to differences in genetic load (*ρ* = 0.04, *p* = 0.48).

**Table 2.** Analyses of deviance.

| *Traits* | *Tests* | *Block* | *G<sub>OR</sub>* | *N<sub>C</sub>* | *G<sub>OR</sub> x N<sub>C</sub>* | *Error dist.* |
| --- | --- | --- | --- | --- | --- | --- |
| $GR_{MP}$ | $\chi^2$ | - | 164*** | - | - | B |
| $\Delta_{growth}$ | $F$ | 1.3<sup>ns</sup> | 14.4*** | 4.9* | 1.3<sup>ns</sup> | N |
| $LT$ | $\chi^2$ | 0.21<sup>ns</sup> | 105*** | 0.7<sup>ns</sup> | 3.9<sup>ns</sup> | NB |
| $FS$ | $\chi^2$ | 10.2* | 212*** | 2.8<sup>ns</sup> | 2.6<sup>ns</sup> | NB |
| $FT$ | $\chi^2$ | 13.7** | 9.2** | 13.4*** | 9.9** | N |
| $N_F$ | $F$ | 4.6<sup>ns</sup> | 4.8<sup>ns</sup> | 162*** | 1<sup>ns</sup> | NB |
| $W$ | $F$ | 0.5<sup>ns</sup> | 20.3*** | 2.6<sup>ns</sup> | 1.1<sup>ns</sup> | N |
| $GR_{SOIL}$ | $\chi^2$ | 30.6*** | 5408*** | 21.5*** | 4.5<sup>ns</sup> | B |
| $Ic$ | $\chi^2$ | - | 36.8*** | 137*** | 0.32<sup>ns</sup> | G |

*Traits*: $GR_{MP}$ : germination rate of mother plants, $\Delta_{growth}$ : vegetative growth rate, $LT$ : life time, $FS$ : flowering start, $FT$ : flowering timespan, $N_F$ : number of fruits, $W$ : seed weight, $GR_{SOIL}$ : progeny germination rate in pots, $Ic$ : competition index. Statistical tests: $\chi^2$ test or F test. Degrees of freedom were 3, 2, 1 and 2, for block effect (Block), geographic origin ( $G_{OR}$ ), number of competitors ( $N_C$ ) and the interaction term between $G_{OR}$ and $N_C$ , respectively. Significance levels are \*\*\*, $p < 0.001$ ; \*\*, $p < 0.01$ ; \*, $p < 0.05$ ; <sup>ns</sup>, $p > 0.05$ . Error dist. is the assumed distribution of error terms: B: Binomial, N: normal, NB: Negative binomial, G: gamma distributions.

#### Phenology

Three phenology-related traits were studied, namely, *lifetime* (*LT*, number of days between plant transplantation and death), *flowering start* (*FS*, number of days between plant transplantation and first flower appearance) and *flowering timespan* (*FT*, number of days between *FS* and plant death). *Lifetime* differed between geographic origins: Asian accessions had the shortest *Lifetime* and European accessions the longest (AS, 53 ± 1 days; ME, 57 ± 1; EU 71 ± 2; AS *vs* ME: Lrt, *χ*^2^ = 4.2, Δ*df* = 1, *p* = 0.045 and ME *vs* EU: Lrt, *χ*^2^ = 46, Δđ*df* = 1, *p* < 0.001; Table 2 and Fig. 2C). The same pattern was observed for *flowering start*: the Asian accessions flowered first and the European accessions last (AS, 23 ± 1 days; ME, 31 ± 1; EU, 44 ± 1; AS *vs* ME: χ^2^ = 33, Δ*df* = 1, *p* < 0.001 and ME *vs* EU: Lrt, *χ*^2^ = 55,Δ*df* = 1, *p* < 0.001; Table 2 and Fig. S2D). Because the interaction term between geographic origin (*G_OR_*) and number of competitor (*N_C_*) was significant for *flowering timespan* (Table 2), a sub-model considering only the block effect and *G_OR_* was also computed. Geographic origin had a weaker, albeit significant, influence on *flowering timespan* than on the two other phenology traits (Lrt, *χ*^2^ = 8.5, *df* = 2, *p* < 0.05). Middle-Eastern and European accessions had the shortest *flowering timespan* (ME, 25 ± 1; EU, 27 ± 1; Lrt, *F* = 2.4, Δ*df* = 1, *p* = 0.12) and Asian accessions the longest (AS, 29 ± 1 days; Lrt, *F* = 5.6, *df* = 1, *p* < 0.02; Fig. S2E1). In addition, genetic load and phenology traits were strongly correlated. Accessions with the highest genetic load had the lowest *lifetime* but also the earliest *flowering start* (respectively, *ρ* = −0.42 and *p* = −0.55; all *p* < 0.001); they also tended to have a longer *flowering timespan* but the correlation was much weaker (*ρ* = 12, *p* = 0.04). Finally, as *flowering start* did not affect *flowering timespan* (*G_LM_, t* = 1.4, *df* = 283, *p* = 0.16), differences in *lifetime* were mainly due to differences in the length of the vegetative period.

#### Reproductive traits

Since the number of flowers and the number of flowering stems were strongly correlated to the total number of fruits (*r* = 0.99 and 0.8, respectively), only the total number of fruits (*N_F_*) was used as a proxy for fertility. Though AS accessions produced the highest number of fruits, no significant variation of fertility was detected between geographic origins (AS, 544 ± 38;ME,475 ± 41; EU,492± 30, Table 2).

### Progeny

#### Seed weight

At the end of the competition experiment, seeds were collected from each plant. The seed weight (W) differed significantly between geographic origins (Fig. S2F and Table 2). AS accessions (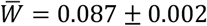 mg/seed) produced lighter seeds than EU (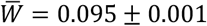; Lrt, *F* = 13.90, *Δd_f_* = 1, *p* < 0.01) and ME accessions (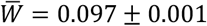; *Lrt, F* = 16.94, *Δdf* = 1, *p* < 0.001), which did not differ significantly between them (*Lrt, F* = 0. 71, *Δdf* = 1, *p* = 0.39). Thus, accessions with the highest proportion of deleterious mutations produced the lightest seeds (*ρ* = −0.23, *p* < 0.001).

#### Germination rate

After having been weighted, seeds were sown in either agar plates or pots to measure progeny germination rate in a favourable and in a more stressful environment, respectively. Results for *GR_AGAR_* and *GR_SOIL_* were similar though, as expected, the differences between factors were stronger for *GR_SOIL_* than for *GR_AGAR_*; only results for *GR_SOIL_* are reported in the main text (results for *GR_AGAR_* are summarized in Table 3). For *GR_SOIL_*, the interaction term between geographic origin (*G_OR_*) and number of competitors (*N_C_*) was significant (Table 2) and a sub model considering only the block effect and *G_OR_* was thus also used. As for mother plants previously, the geographic origin explained a large part of the variance in germination rate (*Lrt, χ*^2^ = 5400, *Δdf* = 2, *p* < 0.001). Seeds from AS accessions had by far the lowest germination rate (only 6.5 %), while 28% and 81% of the seeds from EU and ME, respectively, germinated (Fig. S2G). Interestingly, *GR_soil_* was correlated to seed weight (p = 0.21, *p* < 0.001), i.e. the heavier the seeds, the higher the germination rate; in *C_bp_*, seed weight is thus a good proxy of seed viability.

**Table 3.** Dynamics of germination among the main geographical areas of origin.

| **Contrast** | **Agar** |  | **Soil** |  |
| --- | --- | --- | --- | --- |
|  | *z* | *p* | *z* | *p* |
| Asia vs Europe | 28.05 | < 0.001 | 24.25 | < 0.001 |
| Asia vs Middle East | 29.93 | < 0.001 | 49.21 | < 0.001 |
| Europe vs Middle East | 5.65 | < 0.001 | 52.62 | < 0.001 |

*z*- and *p*-values obtained from Cox's hazard models of germination dynamics for all geographic areas in two different conditions (soil and agar).

#### Complex phenology

In both environments (soil and agar), the *germination time* differed between geographic origins (Cox’s hazard model, all *p* < 0.001; Table 3 and Fig. 3), AS accessions germinating later (*GD_SOIL_* = 6.26) than both EU (*GD_SOIL_* = 4.75) and ME accessions (*GD_SOIL_* = 4.10). However, as for mother plants, progenies (*FS_pro_*) of AS accessions started to flower earlier (~23 days) than ME (~31 days) and EU accessions (~36 days) despite their slightly late germination time. This indicated that AS accessions reached their reproductive phase much faster than accessions from EU or ME. Interestingly, this trait is much more conserved between mother plants and offspring in Asian accessions (*r* = 0.81) than in EU (*r* = 0.46) or ME accessions (*r* = 0.09).

**Figure 3.**
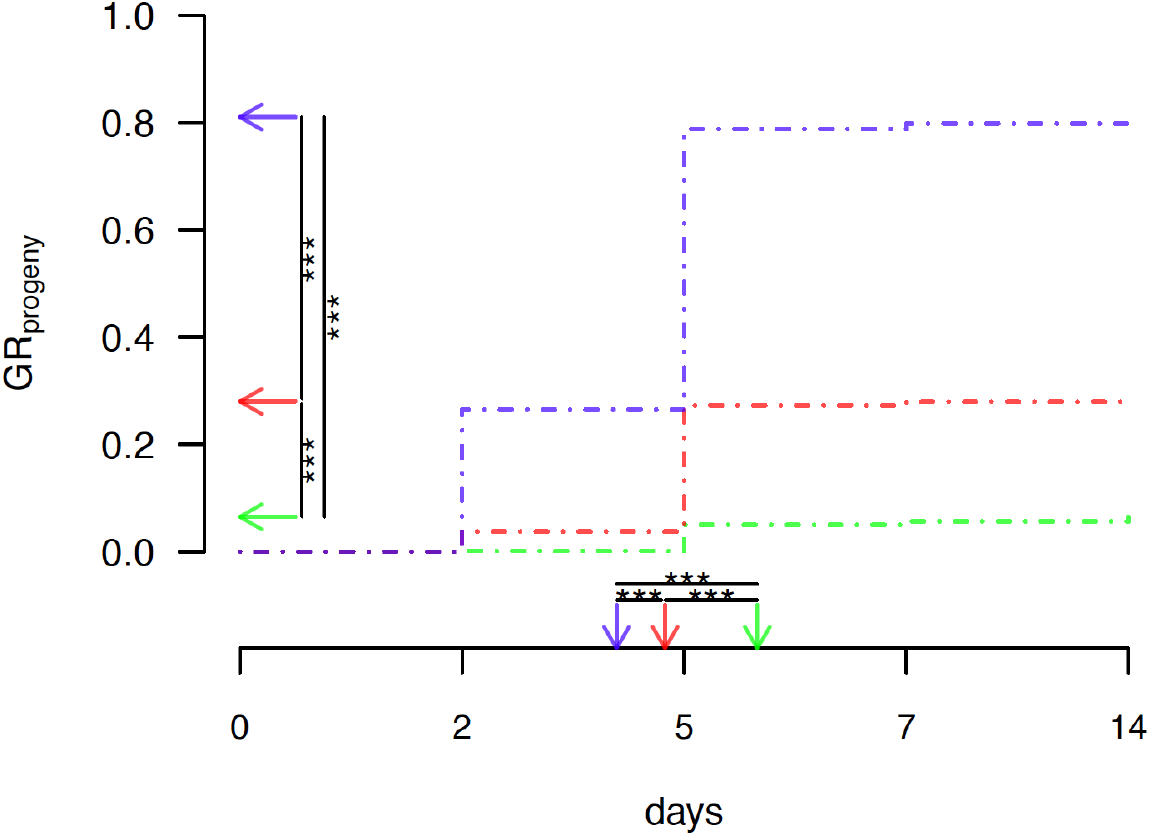
Germination rate of the progeny (*GR_progeny_*) in soil at different time points (days) according the geographical origin. The dashed lines indicate the dynamic of the germination for the three geographical area: AS (green), EU (red), ME (blue). The vertical arrows indicate on the x-axis the mean time to germinate and the horizontal arrows the proportion of germination reach for each geographical area. The stars indicated the significant differences between the three regions (*P* < 0.05 *, *P* < 0.01 **, *P* < 0.001 ***).

In summary, Asian accessions of *C. bursa-pastoris*, presented the worst performances for life history traits covering the whole life-cycle, germination success, vegetative growth, lifetime and seed-weight, which could partly be explained by a higher genetic load. However, these accessions also flowered much earlier and for a longer period than the accessions from Europe or the Middle-East and thus tended to produce more fruits although this difference was not statistically significant. Paradoxically, the *flowering start* was also the trait with the highest correlation with genetic load: the higher the genetic load, the earlier the flowering.

### Unexpectedly, Asian accessions were the least sensitive to competition

To quantify how the presence of competitors affected *C. bursa-pastoris* life history traits, accessions from Asia, Europe and the Middle-East were grown in absence or with 1, 2, 4 or 8 competitors (*Matricaria chamomilla*).

#### Life history traits

The number of competitors (*N_C_*) affected negatively many life history traits across the whole life-cycle (Fig. 2). First, overall *rosette growth rates* tended to decrease when *N_C_* increased, though it should be pointed out that this effect was mainly driven by a strong decrease in rosette size in the presence of eight competitors Table 2 and Fig. S2B2). *Flowering-timespan* was the only phenology-related trait affected by the number of competitors (Table 2) and it decreased as competition intensity increased (*t* = −3.6; *p* < 0.001; Fig. S2E2). While no significant difference in *fertility* (number of fruits, *N_F_*) was observed between geographic areas, fertility was strongly affected by the intensity of competition (*z* = −7.2; *p* < 0.001; Table 2); the larger the number of competitors, the lower the number of fruits produced (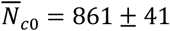, 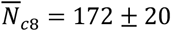; Fig. S2H). Because fertility is one of the main components of fitness, the ability of *C. bursa-pastoris* to withstand competition should strongly determine its fitness. Finally, it is interesting to note that competition intensity did not only affect the mother plant performances but also progeny establishment success as the germination rate of progeny (*GR_SOIL_*) decreased when the mother plant had to face a strong competition (*z* = −2.17; *p* = 0.03; Table 2).

#### Competitive indices

To investigate whether plant ability to withstand competition differed between accessions from different geographic origins (*G_OR_*), a competitive index (*I_cki_*) was computed for each accession *k* and each number of competitors, *i*; the higher the competitive index, the higher the ability to withstand competition (see Methods, eq. 2). As expected, the competitive index decreased when the competition increased but more importantly the magnitude of the effect varied between geographic origins (Table 2 and Fig. 4A). Surprisingly, whatever the number of competitors considered, Asian accessions had higher *I_C_* than Middle-Eastern (*Lrt, χ*^2^ = 5.5, *Δdf* = 2, *p* = 0.001) or European accessions (*Lrt, χ*^2^ = 15.5, *Δdf* = 2, *p* < 0.001). The latter were the most affected by the presence of competitors though not significantly more than Middle-Eastern accessions (*p* = 0.15). In striking contrast with our expectation the correlation between *competitive index* and the genetic load was positive (*ρ* = 0.67, *p* = 0.006, Fig. 4B). This unexpected pattern is explained by the capacity of AS accessions to bloom early, thereby avoiding competition (*ρ* = −0.79, *p* < 0.001, Fig. 4C). To test that hypothesis, a generalized linear model with the *genetic load* and the *flowering start* as explanatory variables was adjusted to the *competitive index* data (gamma error term). The relationship between *genetic load* and *competitive index* was not significant (*Lrt, χ*^2^ = 3.2, *Δdf* = 1, *p* = 0.07) while the negative relationship between *flowering start* and *competitive index* remained highly significant (*Lrt, χ*^2^ = 12.5, *Δdf* = 1, *p* < 0.001).

**Figure 4.**
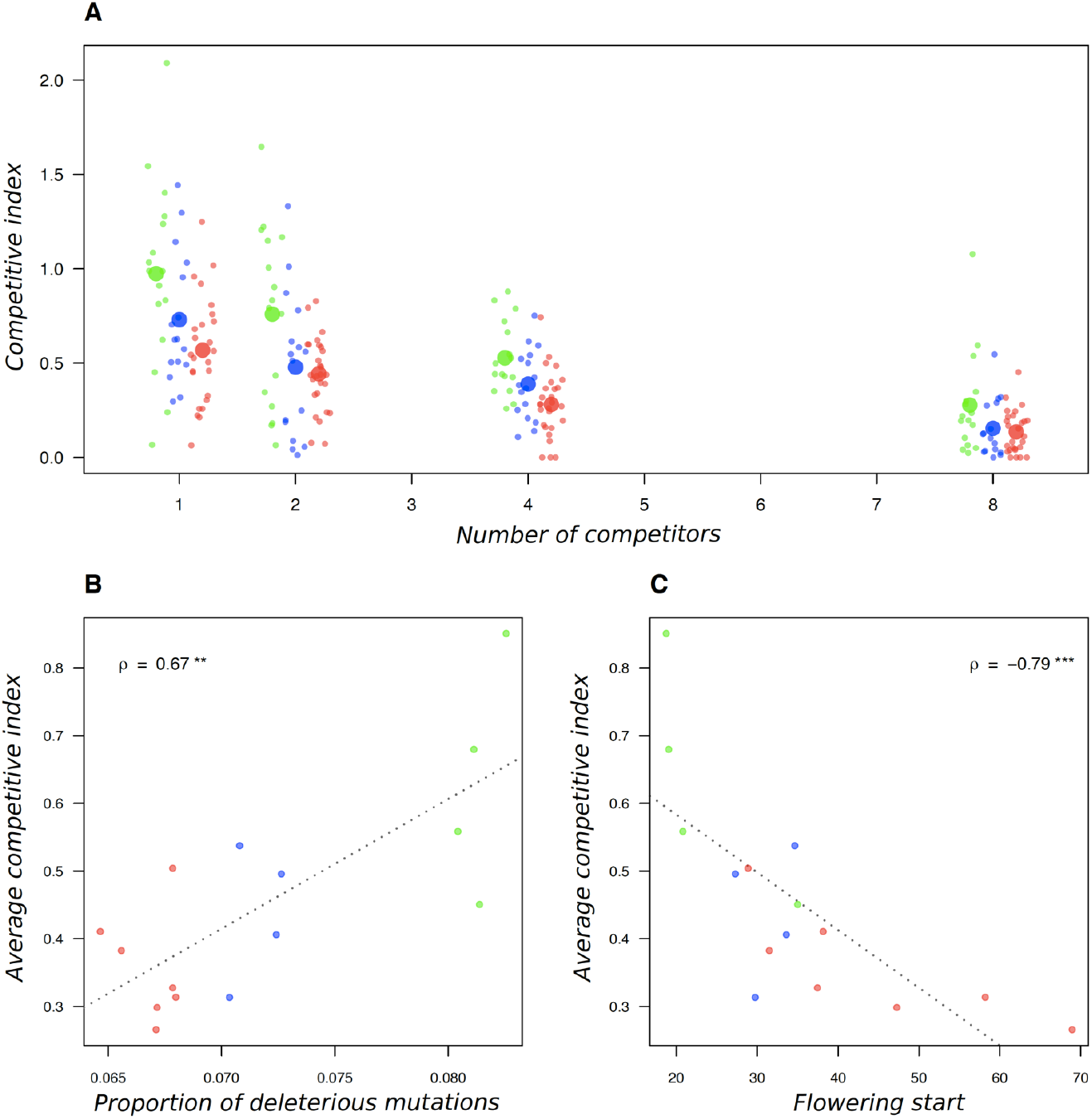
Competition sensibility of *C. bursa-pastoris* accessions. **A-** Competitive index (*I_C_*) as a function of the number of competitors (*N_C_*) for Asian accessions in green, Middle-East accessions in blue and Europeans accessions in red. Large dots are the mean *I_c_* for each geographic origin. **B-** Accession sensitivity to competition (average *I_C_* across the four replicates for each competitor number) as a function of the proportion of deleterious mutations or as a function of of the *flowering start* (**C**). For both relationships, Spearman’s rho correlation coefficients are indicated as well as their significance levels: **, *P* < 0.01; ***, *P* <0.001.

## Discussion

In the present study we analyzed how accessions of *C. bursa-pastoris* from different parts of the natural range performed under increasing density of competitors. The higher the density, the more the focal plant competes for light, acquisition of soil nutrients and water. Here we showed that competitor density had a negative impact on both vegetative growth and reproductive output in term of fruit production for all accessions. However, somewhat unexpectedly given their higher genetic load and their lower absolute performance, Asian accessions were less affected by competition than European and Middle-Eastern ones. A potential explanation could be the shorter time span observed in Asian accessions between germination and first flowering, which would allow limiting the competition for resources acquisition before flowering. Shorter growing period before flowering would explain both the lower absolute performance and the weaker sensitivity to competition. Hence, in spite of their higher genetic load, the ecological strategy of Asian accessions would allow them to cope with competition by avoiding it. However, we don’t know if this efficient strategy in green house conditions also holds in the wild.

Across its natural range, the shepherd’s purse is a ruderal species growing in open environments such as roadsides, wastelands, and more generally any disturbed environment, where competition might not be the main threat. The main aim of the present study was neither to “mimic” natural conditions nor to assess the outcome of competition, but rather to study the response of accessions with different history and levels of genetic loads to varying selection pressures induced by competition (see also similar approach in *Drosophila*, Laffafian et al. 2010; Agrawal & Whitlock 2012; Yun & Agrawal 2014, or for the effect of competition on inbreeding depression, Cheptou et al. 2000). Despite this limitation, the results provide information on the ecological strategy of shepherd’s purse and how it may react in the wild to varying levels of inter-specific competition.

### Possible effect of deleterious mutations throughout the plant life cycle

The mean fitness of individuals in a population can be affected by bottlenecks and admixture events. These demographic events could either increase or decrease the mean fitness and by consequence could have a strong influence on the success of introduction and maintain of the invasive population in new habitats. During a colonization event, it is not rare to observe strong bottlenecks causing decrease in genetic diversity. This can lead to the purging of the strongly deleterious (especially recessive) mutations, thereby increasing mean fitness (Facon et al. 2011) or, on the contrary, the fixation of deleterious mutations (especially weaker ones) leading to a permanent lowering of fitness of the populations at the front of the expansion range (Pujol et al. 2009).

Deleterious mutations can impact reproductive rate and can also have an impact on juvenile competitive ability (Travis et al. 2007; Peischl et al. 2013). Our study validate this expectation because Asian accessions, *i.e*. populations at the limit of the natural range, exhibited a higher proportion of deleterious mutations and also had the lowest performance in term of reproduction, in particular a weak germination rate and a very low progeny viability.

However, although we noticed a decrease in overall fitness in Asian accessions, trade-offs between reproductive traits limited its negative impact. Indeed, the deficiency in germination rate of Asian accessions was partly compensated by the number of fruits produced that tended to be more important than for accessions from the core of the range (AS, 544 ± 38; ME, 475 ± 41; EU, 492 ± 30). Asian seeds were lighter than European and Middle-Eastern ones and likely had a small endosperm and a lack of nutrient reserves. Since resources are not infinite, a trade-off between number of offspring and size of the progeny (e.g. a plant produces a lot of seeds but small in size) is often observed (Stearns 1989). Indeed, in flowering plants, Karrenberg and Suter (2003) showed that a diminution of seed number was linked to an increase in seed mass, which was reinforced by seed longevity (*i.e*. seed survival). Moreover, a trade-off between size and survival of the progeny has also been demonstrated several times in different organisms (Smith et al. 1989; Einum & Fleming 2000). Finally a general tradeoff between offspring number, and offspring quality may be expected, in particular in population on the expansion front as a decrease in seed size (and therefore in resources available for the embryo) could allow for better dispersion (Ganeshaiah & Shaanker 1991).

### Deleterious mutations, phenology, and competition

A good way to compensate for this decrease in mean fitness and to be able to maintain a good colonization ability would be to minimize competition with other species, in particular with a sibling species living in similar conditions, *C. orientalis*. In flowering plant, phenology plays a major role in establishment success in new environments or on colonization fronts. More specifically, the timing of flowering is an important component of the ecology of plants because it plays a crucial role in term of community assembly (Rathcke & Lacey 1985; Sargent & Ackerly 2008). Thus a priority effect, *i.e*. the invasive species bloom earlier than native ones, may contribute to plant communities structure and permit to limit the impact on reproductive success of competition between species for pollinators, nutrient resources, light accessibility (Wolkovich & Cleland 2011). Once again, our results fit well with the hypothesis of priority effect showing that the accessions that came from the front of expansion flowered earlier than those from the core of the range. This phenological shift and the rapid development of Asian accessions permit an easier access to resources (light and nutrients) and limit competitors impact. In agreement with our results, the Asian accessions suffered less from the presence of competitors than the European and Middle-eastern ones. Further experiments with competitors installed at different times could allow us to understand if the lower sensitivity of Asian accessions to competition is due to the shift in phenology limiting the impact of competitors or if some other traits, for example a more developed root system permitting a better access to nutrient resources underground, are responsible for it. It should be noted, however, that our experiment considered only one set of environmental conditions and that we cannot exclude that our results would have been different using different set of parameters. However, a common garden experiment carried out in three localities (Sweden, 59°N; Canada, 43°N; and China, 23°N) with different climatic conditions showed the same pattern (AS before ME and EU) in term of flowering phenology and the same absolute lower performance of Asian accessions (Cornille et al. 2018), suggesting that our results are rather consistent.

So far we have tried to provide an ecological explanation to the difference in response to competition between Asian and accessions from the core of the range. However, other, non-ecological factors may also contribute. First, the shift in phenology may simply be a direct consequence of the higher genetic load of the Asian accessions and may not be an ecological adaptation. Second, introgression from a locally adapted species, *C. orientalis*, in the Asian populations of *C. bursa-pastoris* may also have influenced phenology and facilitated range expansion (Kryvokhyzha et al. 2019). Indeed, some examples in flowering plants tend to show that the introgression with native species could facilitate invasion by an alien species. For instance, hybrids resulting from crossing and introgression between *Carpobrotus edulis* and its native congener *C. chilensis* are known to be successful invaders of coastal plant communities (Albert et al. 1997; Vilà et al. 2000). Another example concerns the annual sunflower, *Helianthus annuus*, which has been able to expand its ecological amplitude and geographic range by introgression with locally adapted native species (Rieseberg et al. 2006). Thus, the introgression of chromosomal segments from a locally adapted species may also have facilitated the expansion of *C. bursa pastoris* in Asia in spite of its higher genetic load.

Numerous studies have detected the presence of negative effects of competition on plant growth and reproduction (Inouye et al. 1980; Walck et al. 1999), but no study, to our knowledge, considered the impact on progeny. However, to understand the impact of competition on individuals with different genetic load, it is important to have an assessment of the overall fitness taking into account traits linking the fitness of the mother plants and the fitness of the progeny. Our study demonstrates that the presence of complex tradeoffs between traits could make populations with high deleterious mutation rate less sensitive to competitors than populations with a lower number of deleterious mutations. A shift in flowering phenology could represent such a tradeoff in the case of populations at the limits of the *C. bursa-pastoris* natural range such as the Asian accessions.

## Acknowledgements

The present study was financed by grants from the Erik Philip Sörensen Foundation and the Swedish Research Council to ML.

## Supporting Information

- **Table S1:** Sampling localities.
- **Table S2:** Correlations with genetic loads.
- **Table S3:** Life history traits measures.
- **Figure S1:** Experimental design.
- **Figure S2:** Effect of geographic origin and number of competitors on life history traits.

