## Supporting information for "Ecological strategy and genetic load in the shepherd’s purse (*Capsella bursa-pastoris*) from the core and the limit of its natural range"

**Table S1.** **Sampling localities.**

| Accession ID | Genetic cluster | Country | Latitude | Longitude |
|---|---|---|---|---|
| DL174 | ASI | China | 38.56 | 121.35 |
| GY37 | ASI | China | 26.37 | 106.43 |
| HJC419 | ASI | China | 30.16 | 120.13 |
| HY85 | ASI | China | 26.53 | 112.33 |
| JZH152 | ASI | China | 30.2 | 112.06 |
| KMB206 | ASI | China | 30.2 | 112.06 |
| N444 | ASI | China | 36.37 | 101.46 |
| NJ219 | ASI | China | 32.03 | 118.46 |
| PL | ASI | China | 23.97 | 120.95 |
| TBS195 | ASI | China | 33.57 | 107.45 |
| TY118 | ASI | China | 37.55 | 112.32 |
| BEL5 | EUR | Russia | 50.55 | 128.28 |
| FR50 | EUR | France | 48.08 | 7.37 |
| HRB132 | EUR | China | 45.45 | 126.37 |
| IRRU2 | EUR | Russia | 52.16 | 104.18 |
| SE14 | EUR | Sweden | 62.64 | 17.94 |
| SE33 | EUR | Sweden | 56.15 | 13.77 |
| STA4 | EUR | United-Kingdom | 56.2 | -2.47 |
| STJ2 | EUR | France | 44.51 | -1.21 |
| VLA3 | EUR | Russia | 43.13 | 131.4 |
| AL87 | ME | Algeria | 35.45 | 7.96 |
| JO56 | ME | Jordan | 31.97 | 35.98 |
| TR73 | ME | Turkey | 41.02 | 28.97 |
| WAC5 | ME | United-States | 31.29 | -97.17 |

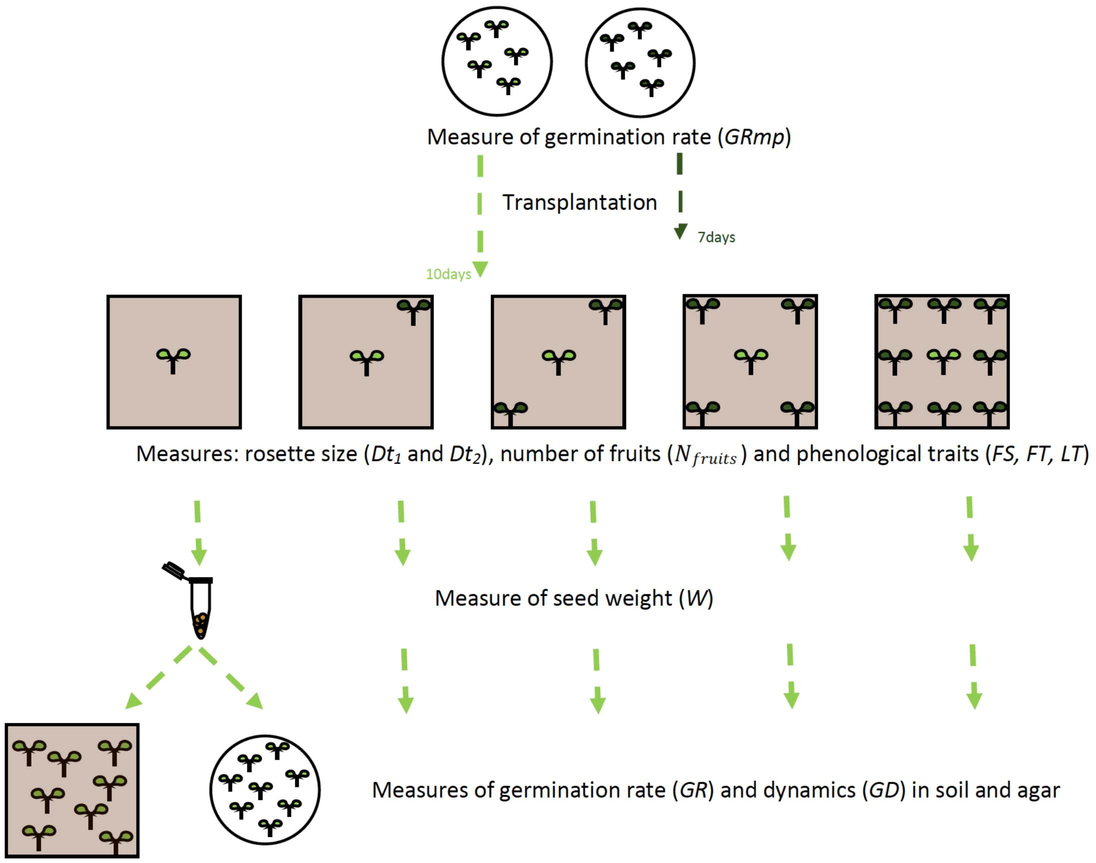


**Figure S1: Experimental design**

**Table S2: Correlation between life history traits and genetic load**

| | Spearman's rho | | | Pearson's product moment | | | |
|---|---|---|---|---|---|---|---|
| | *rho* | *S* | *p* | *r* | *t* | *df* | *p* |
| GRmp | -0.49 | 3417 | 0.02 | -0.46 | -2.4 | 22 | 0.02 |
| DeltaGrowth | 0.04 | 3696900 | 0.482 | -0.03 | -0.5 | 283 | 0.64 |
| LT | -0.42 | 6061300 | <0.001 | -0.38 | -7.1 | 293 | <0.001 |
| FS | -0.55 | 6646500 | <0.001 | -0.52 | -10.4 | 293 | <0.001 |
| FT | 0.12 | 3768800 | 0.041 | 0.13 | 2.3 | 293 | 0.023 |
| SW | -0.23 | 4691200 | <0.001 | -0.25 | -4.4 | 282 | <0.001 |
| $GR_{SOIL}$ | -0.08 | 3996000 | 0.178 | -0.23 | -4.0 | 279 | <0.001 |
| $GR_{AGAR}$ | -0.21 | 4491000 | <0.001 | -0.36 | -6.5 | 279 | <0.001 |

**Table S3: Average trait values per geographic origin and number of competitors**

| | Traits | $N_c$ | Asia | Europe | Middle-East | $\mu \pm sd$ |
|---|---|---|---|---|---|---|
| Vegetative | $\Delta_{growth}$ | 0 | 0.79 ± 0.39 | 0.85 ± 0.38 | 1.21 ± 0.65 | 0.95 ± 0.23 |
| | | 1 | 0.95 ± 0.53 | 0.86 ± 0.42 | 1.6 ± 0.32 | 1.14 ± 0.4 |
| | | 2 | 0.91 ± 0.45 | 0.97 ± 0.38 | 0.88 ± 0.6 | 0.92 ± 0.05 |
| | | 4 | 0.93 ± 0.48 | 0.83 ± 0.34 | 1.33 ± 0.47 | 1.03 ± 0.26 |
| | | 8 | 0.66 ± 0.52 | 0.84 ± 0.43 | 0.93 ± 0.54 | 0.81 ± 0.14 |
| | | $\mu \pm sd$ | 0.85 ± 0.12 | 0.87 ± 0.06 | 1.19 ± 0.3 | - |
| Phenology | *LT* | 0 | 58.31 ± 13.07 | 73.03 ± 20.64 | 58.31 ± 11.44 | 63.22 ± 8.5 |
| | | 1 | 51.56 ± 11.23 | 71.29 ± 18.17 | 54.31 ± 7.12 | 59.05 ± 10.69 |
| | | 2 | 54.06 ± 9.66 | 70.83 ± 16.57 | 54.19 ± 8.57 | 59.69 ± 9.64 |
| | | 4 | 50 ± 10.17 | 66.15 ± 14.79 | 55.62 ± 10.17 | 57.26 ± 8.2 |
| | | 8 | 50.19 ± 6.05 | 70.64 ± 19.65 | 60.69 ± 14.41 | 60.51 ± 10.23 |
| | | $\mu \pm sd$ | 52.82 ± 3.47 | 70.39 ± 2.55 | 56.62 ± 2.81 | - |
| | *FT* | 0 | 33.94 ± 12.06 | 31.73 ± 10.33 | 27.69 ± 9.13 | 31.12 ± 3.17 |
| | | 1 | 27.94 ± 6.24 | 29.79 ± 8.68 | 25 ± 7.95 | 27.58 ± 2.42 |
| | | 2 | 30.31 ± 8.41 | 27.14 ± 8.02 | 22.25 ± 7.17 | 26.57 ± 4.06 |
| | | 4 | 27.94 ± 5.95 | 24.15 ± 6.28 | 24.12 ± 7.09 | 25.4 ± 2.2 |
| | | 8 | 27 ± 7.71 | 22.09 ± 9.24 | 27.44 ± 13.9 | 25.51 ± 2.97 |
| | | $\mu \pm sd$ | 29.43 ± 2.8 | 26.98 ± 3.95 | 25.3 ± 2.3 | - |
| | *FS* | 0 | 24.38 ± 9.02 | 41.3 ± 14.75 | 30.62 ± 4.33 | 32.1 ± 8.56 |
| | | 1 | 23.62 ± 7.74 | 41.5 ± 14.75 | 29.31 ± 3.81 | 31.48 ± 9.13 |
| | | 2 | 23.75 ± 8.09 | 43.69 ± 13.96 | 31.94 ± 7.22 | 33.13 ± 10.02 |
| | | 4 | 22.06 ± 7.08 | 42 ± 12.98 | 31.5 ± 8.08 | 31.85 ± 9.97 |
| | | 8 | 23.19 ± 8.48 | 48.55 ± 21.29 | 33.25 ± 7.88 | 35 ± 12.77 |
| | | $\mu \pm sd$ | 23.4 ± 0.86 | 43.41 ± 3.02 | 31.32 ± 1.47 | - |
| Fertility | $N_F$ | 0 | 786 ± 278 | 906 ± 301 | 891 ± 334 | 861 ± 65 |
| | | 1 | 755 ± 362 | 502 ± 280 | 632 ± 269 | 630 ± 126 |
| | | 2 | 564 ± 326 | 422 ± 245 | 411 ± 350 | 466 ± 86 |
| | | 4 | 411 ± 149 | 320 ± 212 | 347 ± 183 | 359 ± 47 |
| | | 8 | 207 ± 179 | 162 ± 119 | 149 ± 158 | 172 ± 30 |
| | | $\mu \pm sd$ | 544 ± 242 | 462 ± 279 | 486 ± 285 | - |
| Progeny | *W* | 0 | 0.089 ± 0.018 | 0.093 ± 0.013 | 0.094 ± 0.008 | 0.092 ± 0.003 |
| | | 1 | 0.083 ± 0.017 | 0.094 ± 0.012 | 0.097 ± 0.006 | 0.091 ± 0.007 |
| | | 2 | 0.087 ± 0.017 | 0.095 ± 0.015 | 0.098 ± 0.009 | 0.093 ± 0.006 |
| | | 4 | 0.09 ± 0.017 | 0.096 ± 0.012 | 0.099 ± 0.008 | 0.095 ± 0.005 |
| | | 8 | 0.087 ± 0.018 | 0.1 ± 0.024 | 0.097 ± 0.011 | 0.095 ± 0.007 |
| | | $\mu \pm sd$ | 0.087 ± 0.003 | 0.096 ± 0.003 | 0.097 ± 0.002 | - |
| | $GR_{SOIL}$ | 0 | 0.085 ± 0.141 | 0.31 ± 0.325 | 0.857 ± 0.141 | 0.42 ± 0.4 |
| | | 1 | 0.045 ± 0.1 | 0.294 ± 0.314 | 0.804 ± 0.178 | 0.38 ± 0.39 |
| | | 2 | 0.074 ± 0.106 | 0.305 ± 0.322 | 0.759 ± 0.213 | 0.38 ± 0.35 |
| | | 4 | 0.08 ± 0.127 | 0.259 ± 0.291 | 0.821 ± 0.124 | 0.39 ± 0.39 |
| | | 8 | 0.044 ± 0.084 | 0.243 ± 0.284 | 0.807 ± 0.227 | 0.36 ± 0.4 |
| | | $\mu \pm sd$ | 0.07 ± 0.02 | 0.28 ± 0.03 | 0.81 ± 0.04 | - |

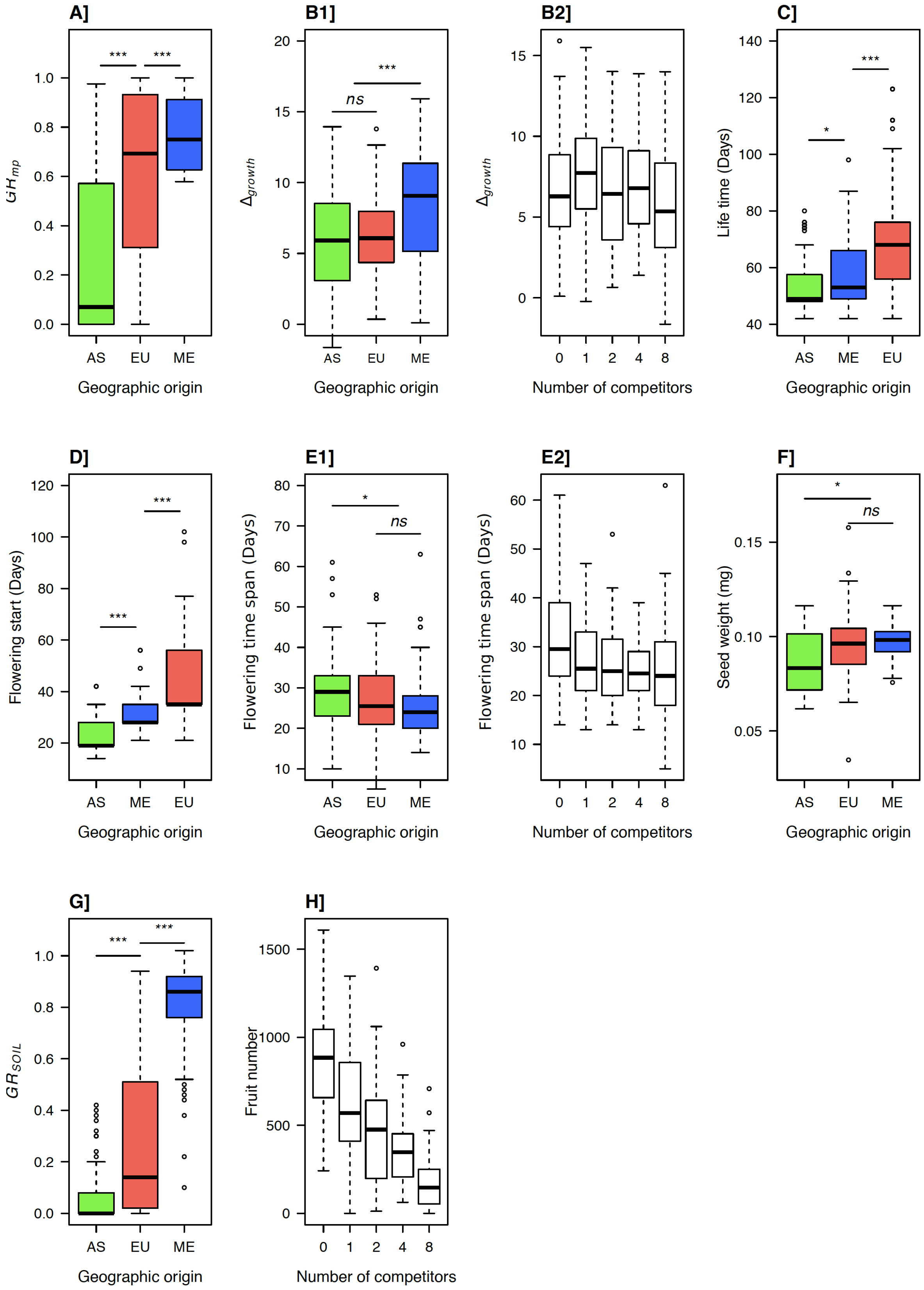


**Figure S2: Effect of geographic origin and number of competitors on life history traits.**
